# Unsupervised clustering and epigenetic classification of single cells

**DOI:** 10.1101/143701

**Authors:** Mahdi Zamanighomi, Zhixiang Lin, Timothy Daley, Xi Chen, Zhana Duren, Alicia Schep, William J Greenleaf, Wing Hung Wong

## Abstract

Characterizing epigenetic heterogeneity at the cellular level is a critical problem in the modern genomics era.
Assays such as single cell ATAC-seq (scATAC-seq) offer an opportunity to interrogate cellular level epigenetic heterogeneity through patterns of variability in open chromatin. However, these assays exhibit technical variability that complicates clear classification and cell type identification in heterogeneous populations. We present *scABC,* an R package for the unsupervised clustering of single cell epigenetic data, to classify scATAC-seq data and discover regions of open chromatin specific to cell identity.

---

Recent advances in single cell technologies such as scATAC-seq^1,2^ and scChIP-seq^3^ have expanded our understanding of epigenetic heterogeneity at the single cell level. However, datasets arising from such technologies are difficult to analyze due to the inherent sparsity. In particular, consider scATAC-seq, designed to interrogate open chromatin in single cells. Open sites in a diploid genome have at most 2 chances to be captured through the assay and only a few thousand distinct reads are generated per cells, resulting in a very low chance that a particular site is captured by the assay. Consequently, it is difficult to determine whether a region is absent in an individual cell due to the lack of openness or due to the sparse nature of data. This creates a challenging task in delineating distinct subpopulations, as only a few genomic regions will have overlapping reads in a large number of cells. To avoid this issue, many studies perform FACS sorting to identify subpopulations, followed by bulk sequencing to determine genomic regions of interest and guide the single cell analysis. If the population is unknown or marker genes are unavailable, then sub-population specific analysis becomes impractical with these techniques.

To combat these challenges and allow for the *de novo* classification of individual cells by their epigenetic signatures, we present a statistical method for the unsupervised clustering of scATAC-seq data, named *scABC* (single cell Accessibility Based Clustering). In contrast to previous works^2,4^ that demand predefined accessible chromatin sites, our procedure relies solely on the patterns of read counts within genomic regions to cluster cells. It requires two inputs: the individual single cell mapped read files and the full set of called peaks (which can be obtained from the union of all of the individual cells without the need for additional experiments). We apply our method to publicly available scATAC-seq data^1, 2, 4^ as well as a true biological mixture to show that our approach can cluster cells with similar epigenetic patterns and identify accessible regions specific to each cluster. We further demonstrate that the cluster specific accessible regions determined by *scABC* have functional meaning and are capable of determining cellular identity. In particular, we show that these cluster specific accessible regions are enriched for transcription factor motifs known to be specific to each subpopulation and that, through association with scRNA-seq data, they can lead to the identification of subpopulation specific gene expression.

## Results

First, we briefly describe our algorithm and the intuition behind it (Fig. 1**a**). To tackle the problem of sparsity, we noted that cells with higher sequencing coverage should be more reliable since important open regions are less likely to be missed by random chance. Therefore, *scABC* first weights cells by (a nonlinear transformation of) the number of distinct reads within peak backgrounds and then applies a weighted *K*-medoids clustering^5^ to partition the cells into distinct groups (see Methods for details). *scABC* uses the ranked peaks in each cell to perform the clustering rather than the raw counts to prevent bias from highly over-represented regions. We found that this usually sufficient to cluster most cells, but a few problematic cells seem to be misclassified. To improve the classification, we calculate landmarks for each cluster. These landmarks depict prototypical cells from each cluster and are characterized by the highest represented peaks in each cluster, which we should trust more than the noisy low-represented peaks. *scABC* finally clusters the cells by assignment to the closest landmark based on the Spearman correlation (Fig. 1**b**). With the cluster assignments we can then test whether each accessible region is specific to a particular cluster, using an empirical Bayes regression based hypothesis testing procedure to obtain peaks specific to each cluster (Fig. 1**c**, Methods).

**Figure 1:**
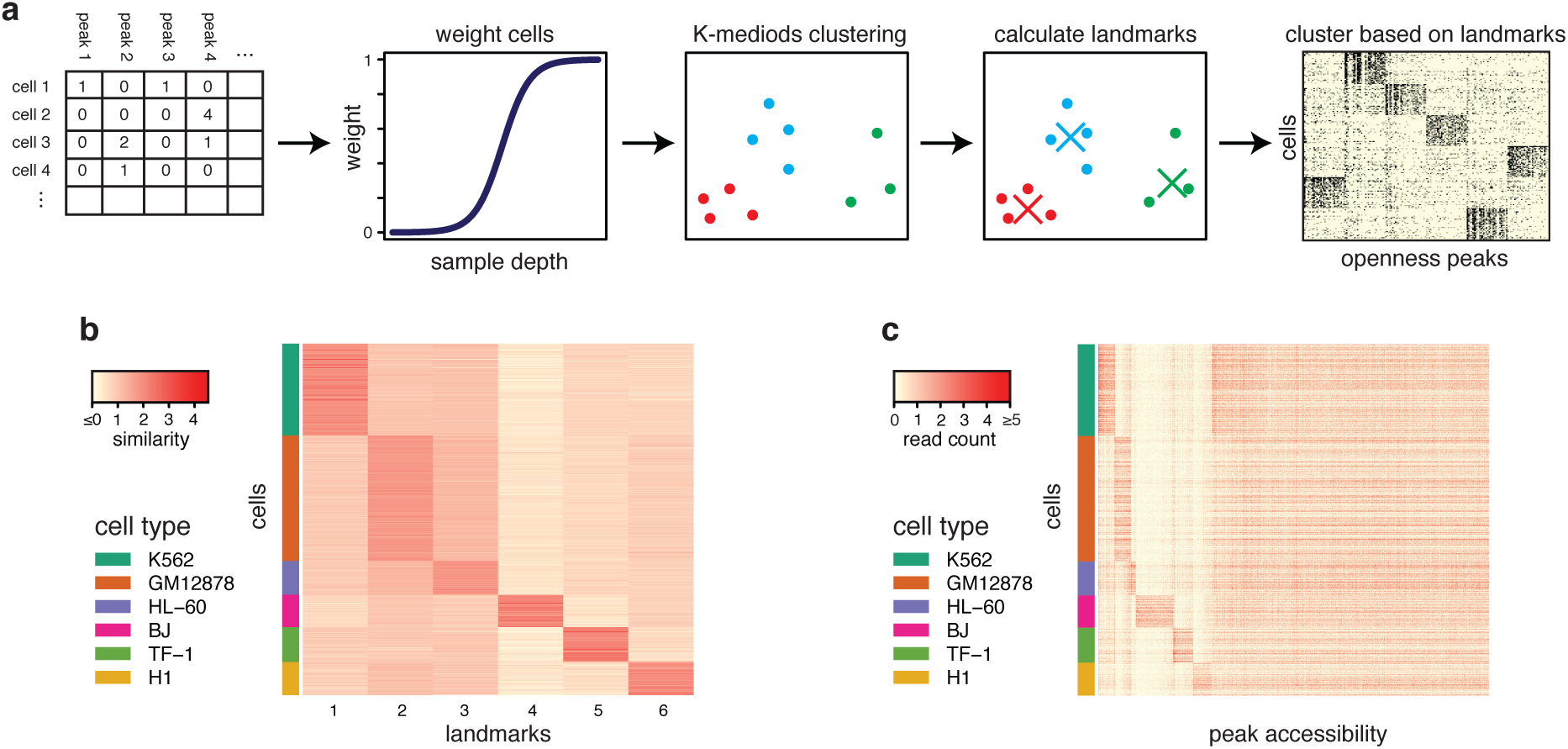
The *scABC* framework for unsupervised clustering of scATAC-seq data. **(a)** Overview of *scABC* pipeline. *scABC* constructs a matrix of read counts over peaks, then weights cells by sample depth and applies a weighted *K*-medoids clustering. The clustering defines a set of *K* landmarks, which are then used to reassign cells to clusters. **(b)** Assignment of cells to landmarks by Spearman correlation, where each cell is highly correlated with just one landmark. The similarity measure used above is defined as the Spearman correlation of cells to landmarks, normalized by the mean of the absolute values across all landmarks for every cell. This allows us to better visualize the relative correlation across all cells. **(c)** Accessibility of peaks across all cells. The vast majority of peaks tend to be either common or cluster specific, allowing us to define cluster specific peaks.

### Performance evaluation using *in silico* mixture of cells

To test our method, we constructed an *in silico* mixture of 966 cells from 6 established cell lines, previously presented in Buenrostro *et al.*^1^ (details in Supplementary Notes, Fig. S1, and Table S1). We then applied *scABC* to this data and determined that there are *K* = 6 clusters using a modified gap statistic (Supplementary Notes, Supplementary Fig. S2). We found 6 well separated landmarks with each cell highly correlated with only one landmark (Fig. 1**b**). The clustering was highly specific with only 4 out of 966 cells misclassified, an error rate of ≈ 0.4% (Supplementary Table S2).

Three major issues are associated with the *in silico* mixture that do not appear in natural mixtures. First, the constructed mixture is inherently biased by batch effects since each cell type must be processed separately. To assess the effect of such bias in our method, we noted that the GM12878 cell line was processed in 4 separate batches, each with the same treatment. We applied *scABC* on the combined 4 batches of GM12878 cells and the results suggested that there is only a single cluster (Supplementary Fig. S2). To further study batch effects, we intentionally set the number of clusters equal to the number of batches. We found that 99% of the cells were associated with two clusters that have similar landmarks and are not dominated by any batches (Supplementary Fig. S3 and Tables S3-S4). We will investigate these two clusters in a later section but these results indicate that *scABC* is robust to batch effects.

The second major issue is that each distinct cell line makes up at least 9% of the *in silico* mixture. We tested how the representation of each sub-population affects discovery by reducing the representation of each cell line in the mixture. We found that some well separated sub-populations, such as BJ and TF1, can be distinguished at 1% of the total population, while other sub-populations such as K562 and HL-60 (both of which are erythroleukemic) may merge when the representation of one falls below 5% of the total population (Supplementary Fig. S4). The last issue is that the *in silico* cell lines are fairly distinct, raising the question: to what extend *scABC* can recognize similar cell types. We designed a test to systematically assess *scABC* sensitivity. For each cell line, we equally divided its cells into two groups and replaced a fraction of peaks in one group using another cell line. Applying *scABC* to these two groups, we achieve successful classifications when at least 50% – 70% of peaks are identical between the groups (Supplementary Fig. S5). In later sections, we will evaluate the sensitivity of *scABC* on real mixtures that have similar sub-populations.

We next investigated whether the cluster specific peaks obtained by *scABC* are able to define cell identity. These peaks contain both narrow and broad regions, as defined by MACS2^6^. In principle, narrow peaks better capture TF binding sites^7^. To measure the enrichment of TF motifs in individual cells we applied chromVAR^8^ to *scABC* defined cluster specific narrow peaks. chromVAR calculates deviations, essentially *z*-scores for TF motif enrichment that are normalized for background accessibility and other biases such as GC content. We found that the most active TFs are typically specific to one or two clusters, identifying active TFs in every cell type (Fig. 2**a**). Some of these TFs were previously shown to be context-specific, for instance, NFKB2 in GM12878 cells^1,2^, SPI1 in HL-60 cells^9^, GATA1::TAL1 in K562 cells^10^, and FOS in BJ cells^11^. It is important to note that TFs with similar DNA-binding motifs show similar motif enrichments. Therefore, POU motifs that are enriched in H1 can demonstrate the activity of POU5F1, the core regulator of human embryonic stem cell self-renewal^12^. We observe that BJ specific TFs seem to be better distinguished than other TFs. Because BJ cells are dissimilar to any of the other cell lines (Fig. 1**b**) and have by far the highest number of cluster specific peaks (Fig. 1**c**), this is not unexpected. We also applied chromVAR to the full set of narrow peaks and found comparable results (Supplementary Fig. S6), indicating that the cluster specific peaks are responsible for the majority of the variation while comprising < 15% of all narrow peaks.

**Figure 2:**
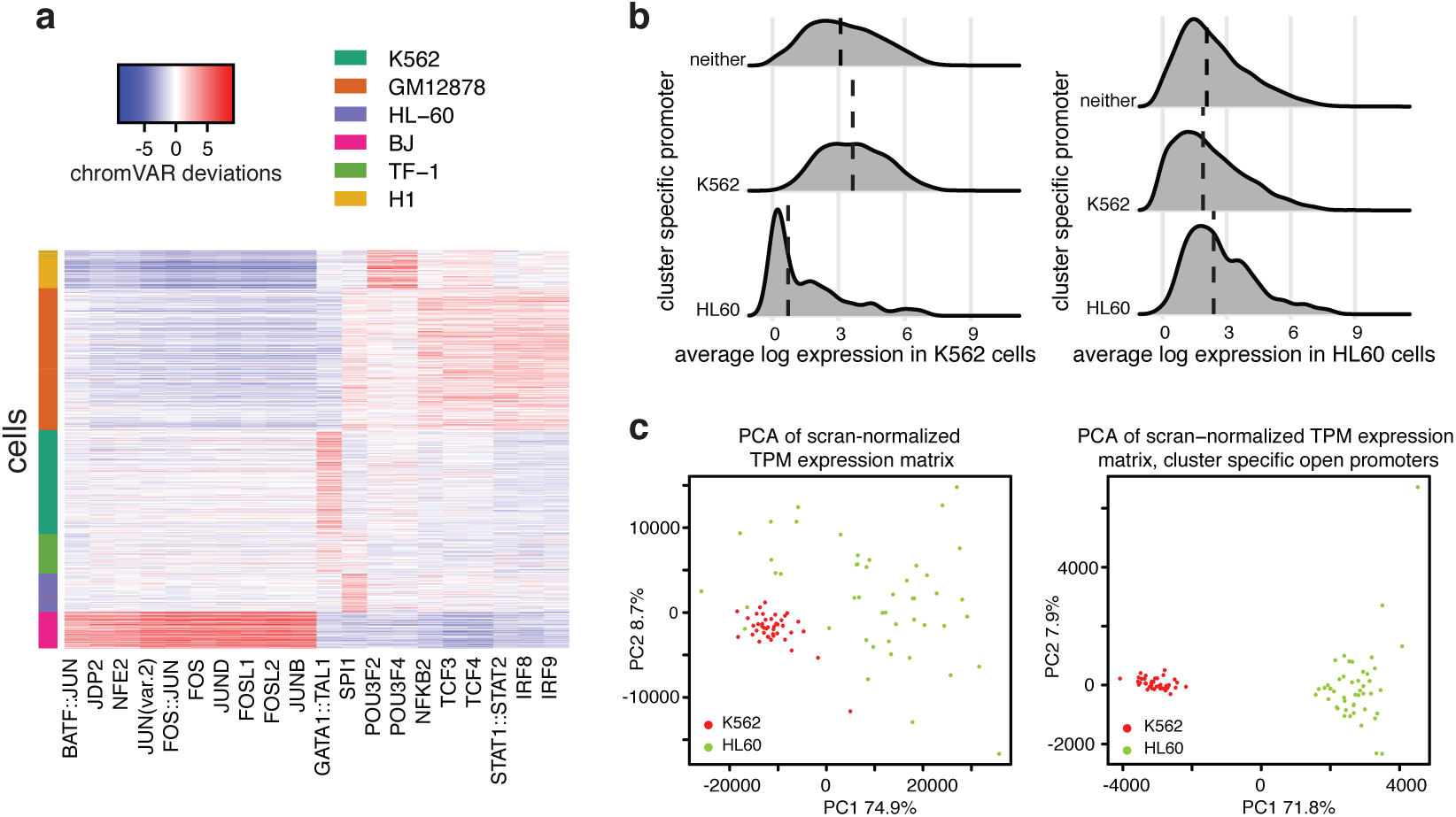
Cluster specific peaks determined by *scABC* shed light on cell identity. **(a)** Application of chromVAR to the cluster specific narrow peaks allows for the identification of cluster specific transcription factor binding motifs. chromVAR calculated deviations are shown for the top twenty most variable transcription factor binding motifs. **(b)** Cluster-specific open promoters distinguish expression. Distribution of the average log gene expression values in genes with either a K562-specific open promoter, HL60-specific open promoter, or non-specific promoter (neither) in K562 cells (left) or HL60 cells (right). **(c)** Integration of scATAC-seq and scRNA-seq enables clear delineation of cell identity. *scABC* applied to scATAC-seq identified genes with cluster specific open promoters for K562 and HL-60 cells. These genes were then used for Principal Component Analysis (PCA) of 42 K562 and 54 HL-60 cells (right) and compared to PCA of all genes (left).

In contrast to narrow peaks, broad regions are more suited to demonstrate functional DNA elements such as promoters and enhancers^13^. We hypothesized that cluster specific broad peaks overlapping gene promoters have functional significance and can help distinguish genes specific to a particular cell type. Specifically, we expect that genes with cell type specific open promoters will have, on average, higher expression in that cell type versus the other cell types in the population^14^. To evaluate this hypothesis, we took 42 K562 and 54 HL-60 deeply sequenced scRNA-seq experiments^15^ (Supplementary Notes). We defined a gene to have a cell type specific open promoter if any open region with an *scABC p*-value of less than 10^−6^ overlapped more than 400 base pairs up to 5 kilobases upstream of the primary FANTOM5 TSS^16^. This cutoff was chosen because it approximately equals the Bonferroni corrected cutoff for a family wise error rate of 0.05.

We first confirmed our hypothesis that genes with cell type specific open promoters tend to be higher in that cell type, compared to other genes (Fig. 2**b**). We next clustered the corresponding gene expression data (in transcripts per million, named TPM) using both all genes and only those genes with cell type specific open promoters in K562 and HL-60 cells (as shown in Fig. 2**c**). After normalization for batch effects^16^, clustering based on all genes did not clearly separate the two cell types in the first two principal components. When genes associated with cell-type specific open promoters were employed, the separation became extremely obvious. Similar patterns were observed when using *t*-SNE plots (Supplementary Fig. S7) This verifies our hypothesis that cluster specific broad peaks shed light on functional significance outside of motif enrichment.

### Performance evaluation on experimental mixtures

In addition to the *in silico* cell line mixture, we examined the capability of *scABC* in classifying three heterogeneous populations. We first applied *scABC* on experimental mixtures of GM12878 and HEK293T cells as well as GM12878 and HL-60 cells^2^. In these experiments, cells were processed in a single batch for each mixture. In both cases, clear separation between the two cell lines were achieved (Supplementary Figs. S8 and S9) that, due to the experimental design, can not be explained by batch effects. Although we correctly classified these cell lines, they are from fairly distinct origins and easy to separate.

To tackle a more difficult problem, we return to the analysis of the GM12878 cell line. Recall that when we intentionally set the number of clusters equal to 4 we found 2 slightly similar clusters. These results were consistent when we set *K* = 2(Supplementary Fig. S10). We hypothesized that these small variations may suggest heterogeneity in the GM12878 cell line. We observed that one cluster is enriched for NF-*κ*B motifs, such as NFKB2, REL, and RELA, and this may be an indication of transcription factor heterogeneity. The nuclear localization of NF-*κ*B was previously shown to dynamically change and cause temporal variations in transcription factor expression^17^, which may explain this heterogeneity. Previous studies^1,2^ have also suggested that cellular variability in GM12878 may be driven by NF-*κ*B heterogeneity. These finding are consistent with our clustering results, but, we cannot further confirm them due to incomplete biological knowledge of GM12878 cell heterogeneity.

### Application to a heterogeneous biological population

For a reliable assessment of our method, we generated a heterogeneous biological population of cells that arise from the same origin. Specifically, we used the hanging drop technique to form embryoid bodies (EBs) from mouse embryonic stem cells (mESCs). We next differentiated EBs using retinoic acid (RA) treatment and performed scATAC-seq on day 4 of the development (Methods). We generated a single 96-well plate and obtained 95 cells that pass quality control (Supplementary Notes).

Our 3D differentiation model (EBs) is expected to produce more interesting heterogeneity than the previous examples. In particular, RA-treated mESCs within 4 days form an outer layer of visceral endoderm cells surrounding ectodermal cells that promote neural properties^18-21^. We therefore hypothesized that the differentiated mESC is a heterogeneous mixture, consisting of both visceral endoderm and neural ectoderm cells. To dissect this heterogeneity in terms of chromatin accessibility, we applied *scABC* to the 95 cells and obtained *K* = 2 clusters (Supplementary Fig. S2) with well separated landmarks (Fig. 3**a**) and cluster specific peaks (Fig. 3**b**). We next ran chromVAR on the cluster specific narrow peaks and found that almost all TFs are specific to one cluster (Fig. 3**c**). The majority of TFs active in cluster 1 play key roles in neural development, including GSX1/2^22^, LBX1^23^, LMX1A^24^, MNX1^25^, NEUROG2^26^, NKX6-1/2^21 27^, UNCX^28^,VAX1^29^, and POU factors^30-32^. The high activity of LHX2/9 in cluster 1 may be related to LHX3/4 (because of their similar motifs), which have been shown to function in the development of mouse motor neurons^33^. In contrast to cluster 1, TFs specific to cluster 2 are essential for visceral endoderm differentiation, such as GATA factors^34^, HNF1A/B^35,36^, and the AP-1 family^37,38^ (i.e. JUN and FOS motifs). The TF enrichment analysis suggests that *scABC* clearly distinguishes neuroectoderm (67 cells) from visceral endoderm (28 cells), two sub-populations with the same origin (mESC) in early embryonic development.

**Figure 3:**
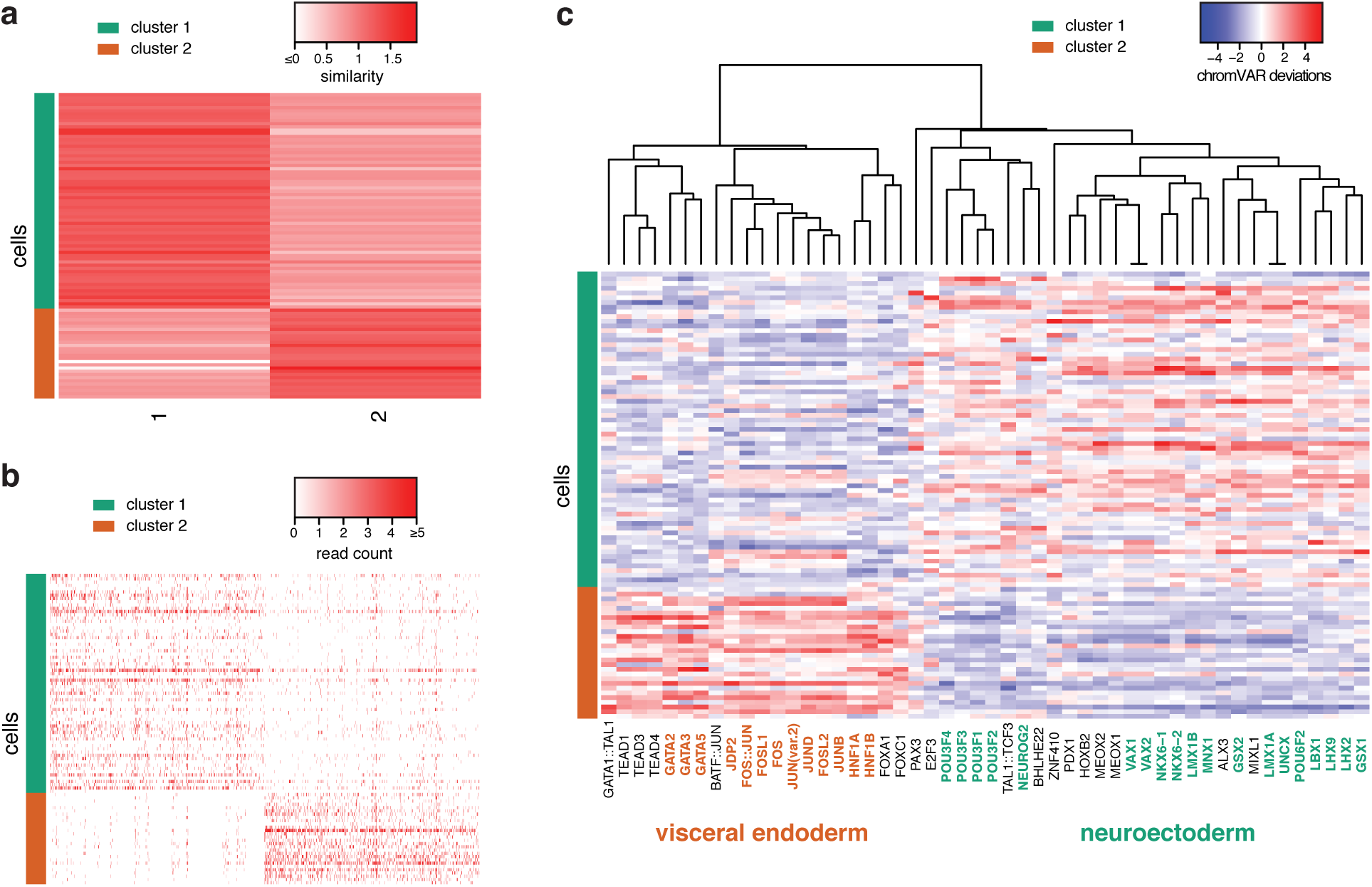
The application of *scABC* to a biological cell mixture. **(a)** 95 scATAC-seq samples were obtained on the day 4 of RA-treated mESC differentiation and classified into two clusters by *scABC.* Here, similarity between cells (rows) and the two detected landmarks (columns) are depicted, with cluster assignments on the left. **(b)** Heatmap for peak accessibility across cluster specific peaks (columns) and cells (rows). To simplify the presentation for each cluster, we only show the top 500 peaks specific to each cluster, i.e. the smallest *scABC p*-values (Methods). **(c)** chromVAR deviations for the top 50 most variable TF motifs (columns) and cells (rows), calculated using cluster specific narrow peaks. Hierarchical cluster analysis of deviations divides motifs into two groups, each specific to just one cluster.

### *scABC* characterizes the leukemic evolution

To further extend the evaluation of *scABC* performance, we tested its ability for detecting developmental stages of cancer evolution. Corces *et al.*^4^ sequenced individual monocytes and lymphoid-primed multipotent progenitors (LMPP) from healthy donors and leukemia stem cells (LSC) and leukemic blast cells (blast) from donors with acute myeloid leukemia. Notably, this dataset is extremely sparse compared to the *in silico* mixture of 6 cell lines (Supplementary Fig. S1). We applied *scABC* followed by chromVAR to the combined mixture of the 390 cells that passed quality control (Supplementary Notes). Our method detected *K* = 2 clusters, which resulted in a clear separation of the cells into a monocyte dominated cluster and a LMPP dominated cluster with blasts predominantly clustered with monocytes and LSCs mainly clustered with LMPPs (Supplementary Fig. S2, Fig. S11, and Table S5, details in Supplementary Notes). When using more clusters, for instance four, the monocyte dominated cluster is stable and well separated from the others but the LMPP is split into two similar clusters (Supplementary Fig. S12 and Table S6). Moreover, one cluster contains only LSCs and blasts, which may be an indication of intermediate stages between LMPP and monocyte. Notably, JUN and JUNB are not enriched in this cluster, and their dysregulation was previously shown to be essential for leukemic stem cell function^39^. In both cases, leukemia cells lie along two major identities on the myeloid progression, represented by monocytes and LMPPs. Our result largely agrees with Corces *et al.*’s study which was based on separate analysis for each of the 4 cell types^4^.

### Comparison with previous methods

*scABC* is the first clustering method specifically designed for scATAC-seq. This required us to compare against simpler methods designed for other types of data. Specifically, we compared *scABC* against simple *K*-mediods using Spearman dissimilarity (without weighting and landmarks) and *K*-means on the log transcripts per million matrix (with the transcript length equal to the peak length) using the read counts in peaks as well as over long intervals (peaks extended to 100kb). To enable a fair comparison, previous methods were applied to the cells that pass *scABC* quality control (Supplementary Notes). We used MACS2 peaks as input for all approaches (Methods and Supplementary Notes). We first applied all methods to the *in silico* mixture of six cell lines. We found that simple *K*-mediods and *K*-means on the the log TPM matrix had a slightly higher misclassification rates (1% for *K*-mediods and 9.6% for *K*-means versus 0.4% for *scABC,* Supplementary Tables S7 and S8), while clustering over long intervals performed noticeably worse (Supplementary Tables S9 and S10). To investigate the performance further, we downsampled each cell line and found that *scABC* is able to identify smaller subpopulations than the other methods (Supplementary Fig. S13). We next applied these clustering methods to the RA-treated EB cells. We found that they either clustered almost all cells into one group (Supplementary Table S11) or the sub-populations identified were not biologically meaningful when we examined TF enrichment (Supplementary Fig. S14, see the previous section for *scABC* clustering results).

To evaluate the performance of *scABC’s* method of determining cluster specific peaks, we used peaks differentially open in the respective bulk data as a gold standard and compared *scABC* to an existing method for identifying differentially expressed genes in single cell RNA-seq, SCDE^40^ (details in Supplementary Notes). We found that the majority of cluster specific peaks identified by *scABC* are differentially open in the respective bulk data and the overlap was much larger than the differentially expressed peaks of SCDE (Supplementary Figs. S15 and S16). We also observed that SCDE calculated cluster specific peaks are not well separated (Supplementary Fig. S17), compared to *scABC* (Supplementary Figs. S8 and S9). We note that since scATAC-seq data tends to be sparser and have lower read counts than scRNA-seq data, it is not surprising that methods developed for scRNA-seq data, such as SCDE, may not easily generalize to scATAC-seq data.

## Discussion

In summary, we developed *scABC* for the unsupervised clustering and identification of cluster specific peaks for single cell epigenetic data. We showed that *scABC* can be applied to scATAC-seq data of complex mixtures to deconvolve the underlying population structure. We should note that in cases where the population cannot be separated into subpopulations, such as when the population lies in a continuum, *scABC* will not be able to separate the population. In our experience, this is usually indicated by a continuously increasing gap statistic. In such cases other tools such as graph embedding^41^ or *k*-mer analysis^8,42^ may be more appropriate.

We showed that the *scABC* identifies informative peaks for downstream analysis. Since *scABC* only uses the read counts within peaks to identify informative peaks, further analysis on the content of the peaks can be done in an unbiased manner while increasing the signal to noise ratio. For example, we showed that *scABC* in conjunction with chromVAR identifies the drivers of cellular heterogeneity in developmental dynamics in the context of retinoic acid induction. In another example, we showed that cell type specific open promoters can better identify cell type specific expression.

*scABC* is available as an open source R package at https://github.com/timydaley/scABC/.

## Methods

### Unsupervised clustering of scATAC-Seq data

The clustering algorithm of *scABC* can be broken down into three steps.

#### Weighted K-medoids clustering

Cells with low sequencing depth are noisy and can negatively impact the clustering result. We implement a weighted version of the *K*-medoids clustering algorithm, where cells with lower sequencing depth are given smaller weight. Let *h_i_* denote a measure of relative sequencing depth for cell *i*, named sample depth (details in Supplementary Notes). The weight for cell *i* is defined as

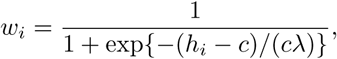

where *c* and *λ* are tuning parameters. As defaults, we use the median of the background and 0.1, respectively. We found that the performance of the clustering is robust to a wide range of {*c, λ*} (Supplementary Table S12).

Let *Y_i_* denote the read counts within peaks for cell *i* (dimension of *Y_i_* is equal to the number of input peaks), *K* the number of clusters, *C* the cluster assignment, and *i_k_* the medoid for cluster *K*, i.e. a cell used as the cluster center. The clustering assignment is given by the solution to

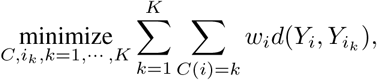

where *d*(·) in general represents the dissimilarly between a pair of samples. We use 1 – Spearman’s rank correlation as the dissimilarity measure, and refer to the Spearman rank correlation as the similarity measure. The problem above is solved by the Partitioning Around Medoids (PAM) algorithm^43^as implemented in the R package *WeightedCluster^5^*.

#### Landmarks

We sum the reads across the cells within a cluster and select the *P* peaks with the highest read counts to obtain the landmark for each cluster identified in the previous step. As a default we set *P* =2,000.

#### Re-clustering using landmarks

To refine the clustering results, we re-cluster the cells by assigning each cell to the landmark with the highest Spearman’s rank correlation using the union of all landmark peaks.

The weighted *K*-medoids algorithm requires the number of clusters *K* in advance. We determine *K* through the gap statistic^44^ with a few modifications to better capture the data structure of single cell experiments, particularly sparsity and cell heterogeneity (Supplementary Fig. S2, details in Supplementary Notes).

### Identification of cluster specific peaks

To find peaks that tend to be more open in one cluster than all others, we formulate the problem in a hypothesis testing framework. We perform the hypothesis testing on all peaks but the procedure is applicable to any subset of peaks, such as narrow or broad peaks. We first introduce our statistical models and then focus on the strategy.

#### Model assumption

Let *K* denote the number of clusters, *R* the total number of peaks, *y_ri_* the read counts for peak *r* in cell *i*, and *x_ik_* the cluster membership for cell *i* with *x_ik_* = 1 if cell *i* belongs to cluster *k* and *x_ik_* = 0 otherwise. We assume that *y_ri_* follows a Poisson distribution with mean *μ_ri_*.

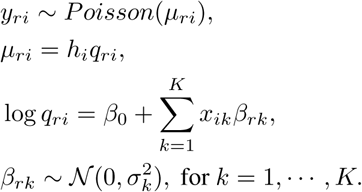

The coefficient *β*_0_ is the intercept and the coefficients *β_rk_* exhibits the effect of the cluster membership on peak *r*. We assume normal priors on the cluster membership effects.

#### Empirical prior estimate

The normal prior enables empirical Bayes shrinkage on *β_rk_*, and stabilizes the noisy estimate when the read counts are low^45^. To obtain a robust empirical prior estimate 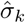, we adopt the quantile matching method proposed in DESeq2^45^. In particular, we first fit a model without the intercept *β*_0_ and without the normal prior to attain the maximum likelihood estimate (MLE) *β^mle^*. Let 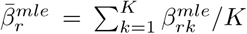; let 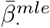 denote the vector 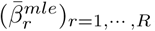; let 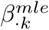 indicate the vector 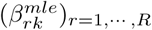; let 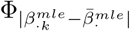 be the empirical cdf of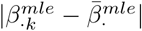, with 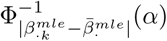 equal to the 1 − *α* quantile of the empirical cdf; and let*Z_α_* be the 1 − *α* standard normal quantile.

The empirical prior estimate for the standard deviation is calculated as

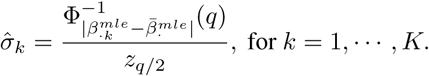

We set *q* = 0.05 in practice. Details for computing *β^mle^* are described in the Supplementary Notes.

#### Hypothesis testing

Suppose Γ_−K_ = {1, …,*K*}_−*k*_ represent the set {1, …,*K*} except for the *k*th element. To test whether peak *r* is specific to cluster *k*, we consider

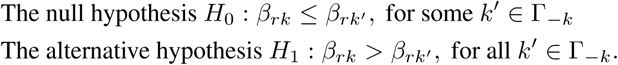

Following the intersection-union test^46^, the null hypothesis can be broken into *K* – 1 simpler null hypotheses *H*_0*k′*_ : *β_rk_*≤*β_rk_’*, with *k′* ∈Γ_−*k*_. For each null hypothesis, the Wald test statistics is, where 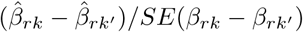 where 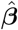 is the maximum a posteriori (MAP) estimate for *β* and *SE*(·) the MAP estimated standard error, which depends on both the observed data and prior estimates. The rejection region for *H*_0*k′*_ with size *α* is

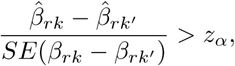

and the rejection region for *H*_0_ with level *α* is

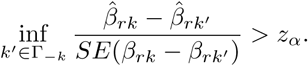

Details for computing 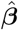 and the standard errors are illustrated in the Supplementary Notes. We finally compute the *p*-value for *H*_0_ as max{*p_k′_*,*k′* ∈Γ_−*k*_}, with *p_k′_* indicating the *p*-value for *H*_0*k’*_.

### Experimental design of RA-treated mESC differentiation

#### Cell culture

Mouse ES cell lines R1 were obtained from ATCC. The mESCs were first expanded on an MEF feeder layer previously irradiated. Then, subculturing was carried out on 0.1% bovine gelatin-coated tissue culture plates. Cells were propagated in mESC medium consisting of Knockout DMEM supplemented with 15% Knockout Serum Replacement, 100 *μ*M nonessential amino acids, 0.5 mM beta-mercaptoethanol, 2 mM GlutaMax, and 100 U/mL Penicillin-Streptomycin with the addition of 1,000 U/mL of LIF (ESGRO, Millipore).

#### Cell differentiation

mESCs were differentiated using the hanging drop method^47^. Trypsinized cells were suspended in differentiation medium (mESC medium without LIF) to a concentration of 37,500 cells/ml. 20 *μ*l drops ( 750 cells) were then placed on the lid of a bacterial plate and the lid was upside down. After 48 h incubation, EBs formed at the bottom of the drops were collected and grown in the well of a 6-well ultra-low attachment plate with fresh differentiation medium containing 0.5 *μ*M RA for 4 days, with the medium being changed daily.

#### scATAC-seq

We followed the scATAC-seq protocol published by Buenrostro et al.^1^ with the following modifications. The EBs were first treated with StemPro Accutase Cell Dissociation Reagent (Thermo Fisher) at 37°C for 10-15 min, followed by vigorous pipetting for another 10 min. The cells were passed through 20 *μ*M cell strainer (pluriSelect) to remove un-dissociated EBs. Before loading, the cells were washed 3 times in C1 DNA Seq Cell Wash Buffer (Fluidigm). 9 *μ*L cells at a concentration of 400 cells/*μ*L were combined with C1 Cell Suspension Reagent at a ratio of 3:2 and 10 *μ*L of this cell mix was loaded on to the 10-17 *μ*M Fluidigm IFC. Single cells were captured using the “ATACseq: Cell Load and Stain (1861×/1862×/1863×)” scripts. After cell capture, IFC was transferred to a Leica CTR 6000 microscope for imaging, followed by Tn5 transposition and primary 8 cycles of PCR using the “ATACseq: Sample Prep (1861×/1862×/1863×)” scripts. The entire volume ( 3.5-5 *μ*L) of the amplified transposed DNA was transferred to a 96-wll plate containing 10 *μ*L of C1 DNA Dilution Reagent. In the 96-well plate, harvested libraries were further amplified in 50 *μ*L PCR (1.25 *μ*M custom Nextera dual-index PCR primers in 1x NEBNext High-Fidelity PCR Master Mix) using the following PCR conditions: 72°C for 5 min; 98°C for 30 s; and total 14 cycles of: 98°C for 10 s, 72°C for 30 s, and 72°C for 1 min. The PCR products were pooled together (4.8 mL) and the pooled library was purified on a single MinElute PCR purification column (Qiagen) and eluted in 20 *μ*L of Elution Buffer. Libraries were quantified using qPCR prior to sequencing using Illumina NextSeq 500 (paired-end 75 bps).

## Acknowledgments

This work was supported by grants R01HG007834, P50HG007735, and R01GM109836 from the National Institutes of Health (NIH). The authors would like to thank Dr. Michael Snyder and members of Wong, Greenleaf, and Chang Labs for helpful suggestions.

## Author Contribution

M.Z, Z.L, T.D, and W.H.W conceived the project. M.Z, Z.L, T.D, Z.D, and A.N.S performed the analysis with inputs from W.J.G and W.H.W. X.C. generated the RA induction data. M.Z, Z.L, and T.D prepared *scABC* R package. All authors wrote the manuscript.

